# The *Plasmodium falciparum* CCCH zinc finger protein ZNF4 plays an important role in gametocyte exflagellation through the regulation of male gametocyte enriched transcripts

**DOI:** 10.1101/2022.03.08.483571

**Authors:** Borja Hanhsen, Afia Farrukh, Gabriele Pradel, Che Julius Ngwa

## Abstract

CCCH zinc finger proteins (ZFPs) function mainly as RNA-binding proteins (RBPs) where they play a central role in the mRNA metabolism. Over 27 CCCH-ZFPs are encoded in the genome of the human malaria parasite *Plasmodium falciparum*, the causative agent of malaria tropica. However, little is known about their functions. In this study, we characterize one member of the PfCCCH-ZFP named ZNF4. We show that ZNF4 is highly expressed in mature gametocytes where it predominantly localizes to the cytoplasm. Targeted gene disruption of *ZNF4* showed no significant effect in asexual blood stage replication and gametocyte development while male gametocyte exflagellation was significantly impaired leading to reduced malaria transmission in the mosquito. Comparative transcriptomics between wildtype (WT) and the ZNF4-deficient line (ZNF4-KO) demonstrated the de-regulation of about 473 genes (274-upregulated and 199 down-regulated) in mature gametocytes. Most of the down-regulated genes show peak expression in mature gametocyte with male enriched genes associated to the axonemal dynein complex formation and cell projection organization highly affected, pointing to the phenotype in male gametocyte exflagellation. Up-regulated genes are associated to ATP synthesis. Our combine data therefore indicate that ZNF4 is a CCCH zinc finger protein which plays an important role in male gametocyte exflagellation through the regulation of male gametocyte enriched genes.

**Author Summary:** CCCH ZFPs have gained significant interest due to their ability to interact with RNA and control different RNA metabolic processes. The role of CCCH ZFPs in the malaria parasite *Plasmodium falciparum* has not been well studied. In this study we report the functional characterization of a PfCCCH-ZFP named ZNF4. Using mouse anti-ZNF4 antisera and a ZNF4-HA tag parasite line we show that the protein is mainly expressed in the cytosol of mature gametocytes. To determine the function of ZNF4, we generated a ZNF4 knockout parasite line and we found that the asexual blood stages and gametocytes developed normally while the ability of the male gametocytes to exflagellate as well as the further development of the parasite in the mosquito was significantly impaired. Interestingly, when the transcriptome of mature gametocytes of the ZNF4-KO was compared to the WT, down-regulated genes were mainly male gametocyte enriched genes associated to processes involved in gametocyte exflagellation such axonemal dynein complex formation and cell projection organization indicating that ZNF4 plays an important role in male gametocyte exflagellation through the regulation of male gametocyte enriched transcripts.

## Introduction

Malaria is one of the deadliest parasitic diseases which resulted in over 241 million infections and 627 000 deaths in 2020 [1]. The disease is caused by apicomplexan parasites of the genus *Plasmodium* with *P. falciparum* as the caustic agent of malaria tropica causing the most severe form. Malaria is transmitted from the human to the anopheline mosquito by a subset of specialized cells, the gametocytes which leave the asexual replication cycle and differentiate into male and female forms. Once mature, the gametocytes are picked up by a blood feeding mosquito. In the midgut of the mosquito, they become activated by external stimuli then the male gametocytes undergo exflagellation, a process that includes three rounds of DNA replication followed by the release of 8 motile microgametes, while the females develop into macrogametes. Following fusion of a microgamete and a macrogamete, a zygote forms during the first hour post-activation, which transforms into an infective ookinete within the following 24 h. The motile ookinete traverses the midgut epithelium before settling down and forming an oocyst between epithelium and basal lamina [2,3].

Gametocyte development as well activation in the mosquito midgut is supported by a well-coordinated sequences of gene activation and silencing events, which are essential to prepare the parasite for transmission from the human to the insect host. Research over the past decade has demonstrated a pivotal role of transcriptional and translational regulation in this process. Gametocyte commitment, the process by which asexual blood stage parasites enter the sexual pathway to form gametocytes, has been shown to be regulated by the transcription factor AP2-G. The expression of AP2-G promotes the transcription of early gametocyte genes, which leads to gametocyte commitment and formation [4–6]. In a recent study, it was shown that another transcription factor AP2-G5 is essential for gametocyte maturation through the down-regulation of AP2-G and a set of genes activated by AP2-G prior to gametocyte development [7]. Other transcription factors such as AP2-FG and AP2-O3 have been shown to regulate gene expression in female gametocytes [8,9] and ookinete development by AP2-O family [10,11].

An important mechanism of transcript regulation is translational repression which has been shown to play an important role in the regulation of female gametocyte transcripts of the malaria parasite. Transcripts of parasite required for mosquito midgut stage formation is synthesized and stored in granules in female gametocytes, where they are translationally repressed by binding to regulatory ribonucleoprotein complexes and the repression is only released after gametocyte activation to promote zygote to ookinete formation [12,13]. In *P. berghei*, the RNA helicase DOZI (development of zygote inhibited) and the Sm-like factor CITH (homolog of worm CAR-I and fly Trailer Hitch) play central roles in the formation of the ribonucleoprotein complex which stores several transcripts including P25 and P28 which are later released after gametocyte activation for zygote to ookinete development [14,15]. In *P. falciparum* on the other hand, the Pumilio/Fem-3 binding factor (Puf) family Puf2 which is an RBP together with its interaction partner 7-helix-1 have been shown to be involved in translational repression of a number of gametocyte transcripts including Pfs25 and Pfs28 [13,16]. Also, the RBP Puf1 has be shown to play an important role in the differentiation and maintenance of mainly female gametocytes [17].

To date, little is known how the transcripts are regulated in male gametocytes. In previous studies, we carried out chemical loss of function studies using inhibitors targeting histone modification enzymes and we showed significant de-regulation of genes expression in immature, mature and activated gametocytes following treatment with the inhibitors [18,19]. These studies indicated epigenetic gene regulation mechanisms during gametocyte development and potentially in male and female gametocytes. In one of the study, we identified a CCCH-ZFP which we named ZNF4 (PF3D7_1134600) that was highly de-regulated following the treatment of the immature gametocytes with the histone deacetylase inhibitor Trichostatin A [18]. CCCH-type zinc finger proteins mainly act as RBP with important roles in the RNA metabolism including RNA stability and transcriptional repression [20,21]. In some cases, CCCH-ZFPs may traffic between the nucleus and the cytoplasm [22] and bind both DNA and RNA [23], thereby functioning in both DNA and RNA regulation [24]. We now show that ZNF4 is a potential nucleic acid-binding protein which plays an important role in microgamete exflagellation through the regulation of male-specific genes during gametocyte development.

## Results

### ZNF4 is a CCCH-ZFP expressed mainly in gametocytes of *P. falciparum*

Analysis of the ZNF4 protein features using UniProt shows a 204 kDa protein with three CCCH zinc finger domains between amino acid 513 to 540, 548 to 574 and 582 to 610 (Fig 1A). In addition, the 3D-structure of the protein as predicted using AlphaFold [25,26] shows the arrangement of alpha helices and the beta strands in the CCCH domains to coordinate the zinc binding (Fig 1B). To determine the transcript expression of ZNF4, a semi quantitative RT-PCR was performed using asexual blood stages (rings, trophozoites and schizonts) as well as gametocyte stages (immature, mature and 30 min post-activated gametocytes). The transcript levels of Pfama1 (apical membrane antigen 1) and Pfccp2 (LCCL domain-containing protein) were determined to control for purity in asexual blood stage and gametocyte samples respectively. Samples without reverse transcriptase (-RT) were used as controls to verify the absence of genomic DNA. Pffbpa (fructose-bisphosphate aldolase) was used as housekeeping loading control. High transcript expression of ZNF4 was observed in the gametocyte stages as compared to the asexual blood stages (Fig 1C). To determine the ZNF4 protein expression, a recombinant peptide (RP) corresponding to a portion of ZNF4 (Fig 1A) was expressed in *E. coli* and used to generate mouse polyclonal antisera against ZNF4. Immunofluorescence assays (IFAs) show mainly a cytoplasmic expression of ZNF4 in both asexual blood stages and gametocytes with the highest expression in mature gametocytes (Fig 1D). Differences in sex specific expression between male and female gametocytes were not observed. To confirm ZNF4 expression, we used a pSLI-ZNF4-HA-glmS parasite line in which the 3’end of the endogenous gene was fused with the sequence of a 3x HA-tag and a glmS ribozyme (Fig S1A). Successful integration in the parasite line was obtained (Fig S1B) and we were able to detect the HA-tagged ZNF4 in mature gametocyte lysates at a molecular weight of roughly 250 kDa (Fig S1C). IFAs using anti-HA confirmed expression of the protein in the asexual blood stages and gametocytes with the highest expression in mature gametocytes (Fig S1D).

**Fig 1:**
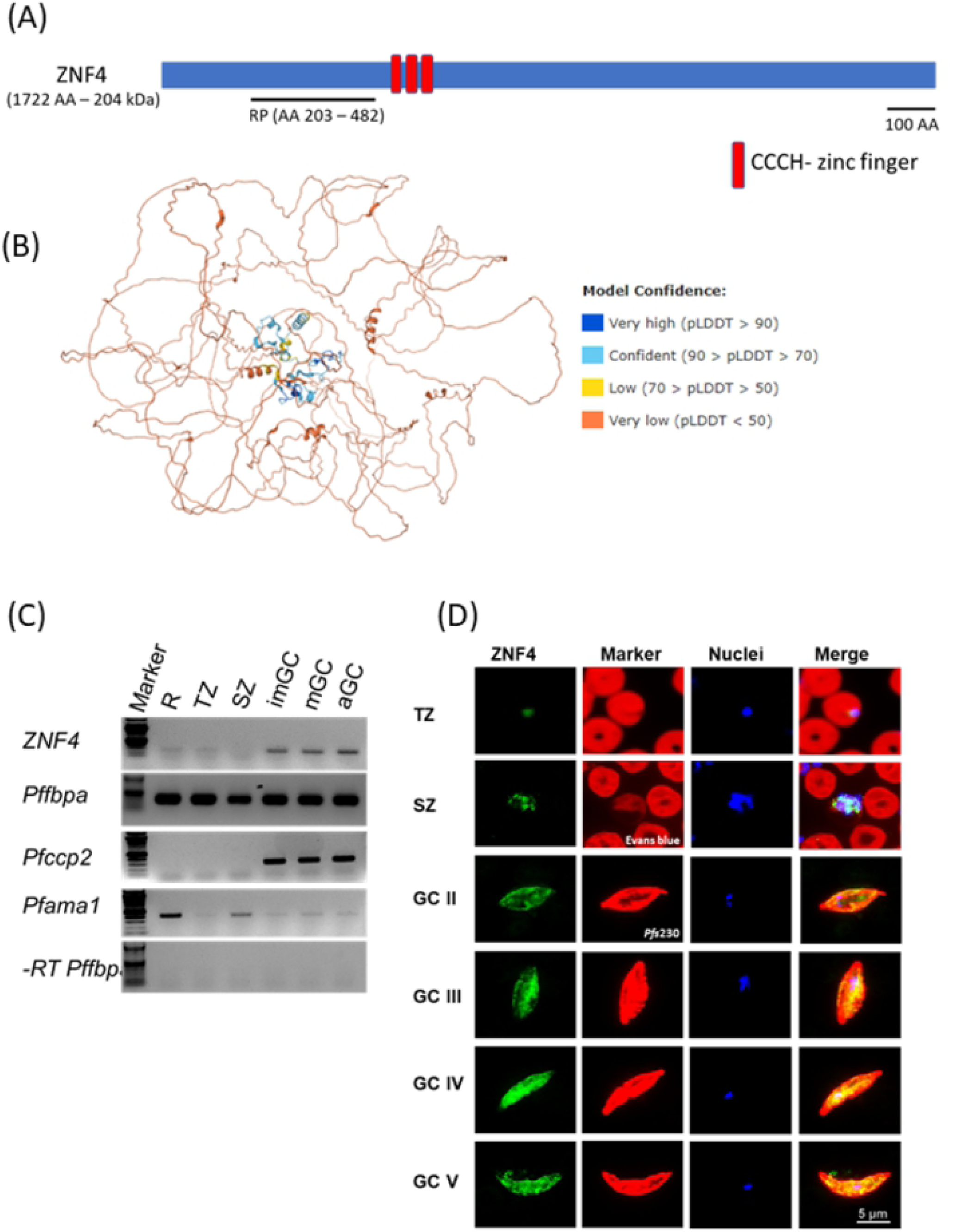
Domain architecture and protein expression of ZNF4. (A) Schematic of the ZNF4 domain structure. The 204 kDa protein contains three CCCH zinc finger domains indicated in red. The black line indicates the region used for the generation of the recombinant peptide. (B) Predicted 3D structure of ZNF4. The 3D structure of the protein was predicted using AlphaFold [25,26]. Colours indicate the confidence level of prediction which ranged from blue (very high confidence) to orange (very low confidence). (C) Transcript expression of ZNF4 in the blood stages of *P. falciparum*. Complementary DNA was synthesized from rings (R), trophozoites (TZ), schizonts (SZ), immature (imGC), mature (mGC) and gametocytes at 30 min post-activation (aGC) and subjected to diagnostic PCR using ZNF4-specific primers. The expression levels of Pfama1 and Pfccp2 were used to verify asexual blood stage and gametocyte-specific expression. Samples lacking reverse transcriptase (-RT) were used as controls to check for any contamination with genomic DNA. Pffbpa was used as a loading control. (D) Localization of ZNF4 in different blood stages of *P. falciparum*. Anti-ZNF4 was used to immunolabel fixed samples of trophozoites, schizonts and gametocytes (GC stage II to V) as well as of activated gametocytes (aGC) at 30 min post-activation (green). Asexual blood stages (trophozoites and schizonts) were visualized by labelling with Evans blue and gametocytes were visualized by using rabbit anti-Pfs230 (red); nuclei were highlighted by Hoechst nuclear stain 33342 (blue). Bar, 5 µm.

### Targeted gene disruption of *ZNF4* does not impact asexual blood stage replication and gametocyte development

To determine the function of ZNF4, we utilized the selected linked integration-mediated targeted gene disruption method [27] to generate a disrupted ZNF4 parasite line, named ZNF4-KO (Fig 2A). These parasites express only a truncated N-terminal GFP tagged fragment which lacks the three CCCH zinc finger domains. Successful disruption was confirmed by diagnostic PCR, which indicated 5’and 3’ integration and lack of wildtype in the ZNF4-KO parasite line (Fig 2B). To further confirm the gene disruption, the remaining part of the ZNF4-KO truncated protein which is fused to GFP was detected by Western Blotting using different parasite stages (rings, trophozoites, schizonts, immature and mature gametocytes) as well as by live imaging (Fig 2C, D). The western blotting with the ZNF4-KO parasite line further confirmed the expression of the truncated protein at the expected molecular weight of 53 kDa in different *P. falciparum* stages (Fig 2C).

**Fig 2:**
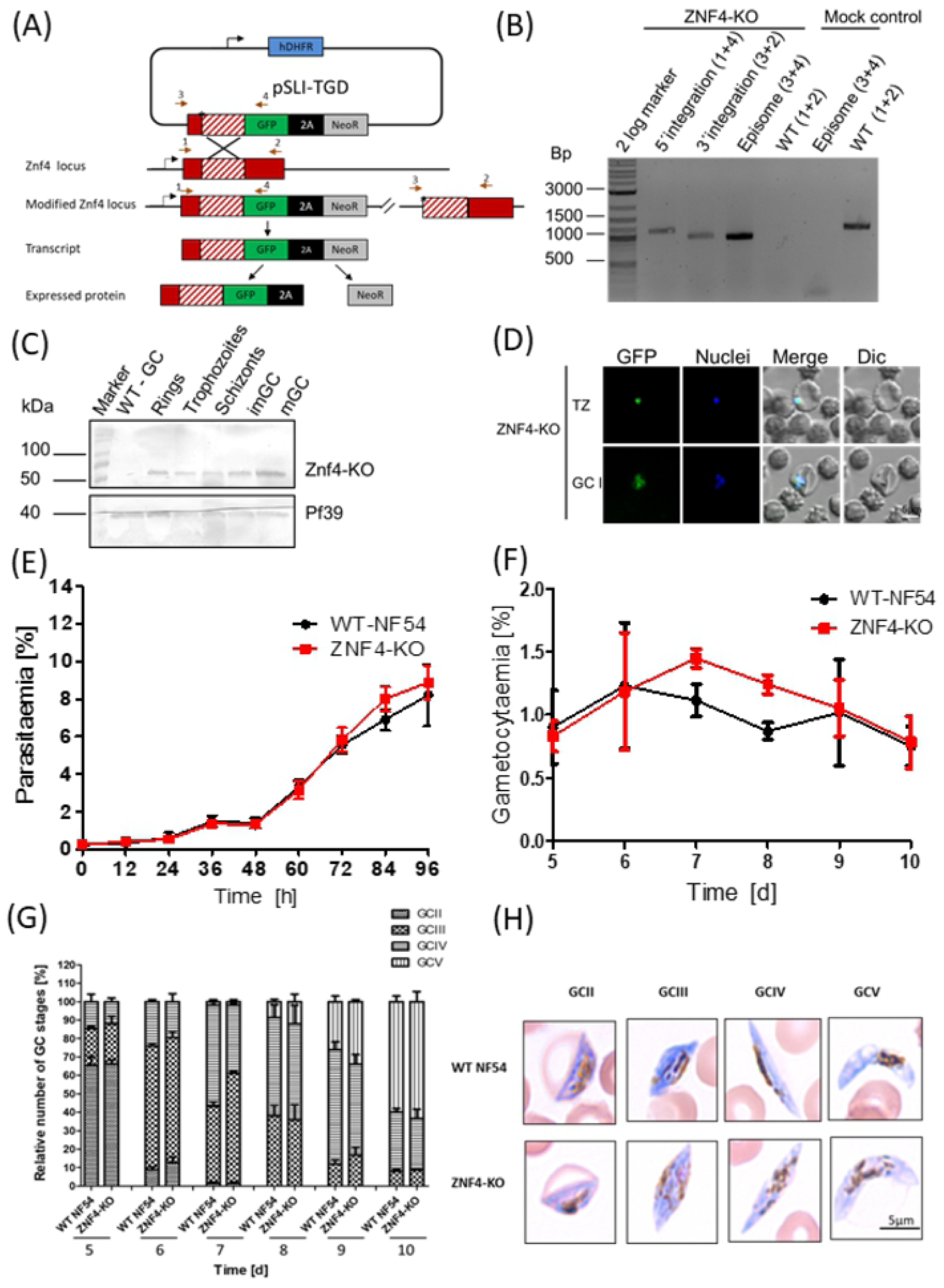
Targeted gene disruption of *ZNF4* and its effect on asexual blood stage replication and gametocytogenesis. (A) Schematic depicting the generation of ZNF4-KO via single crossover recombination-based gene disruption using selective linked integration targeted gene disruption (SLI-TGD). The vector pSLI-TGD was modified to contain a 601 bp sequence block (white box with red stripes) from near the 5’ end of the *ZNF4* coding region (red box). The coding region was maintained in frame with a GFP protein coding region (green box), a 2A “skip” peptide (black box) and the Neo-R gene (grey) that provides resistance to the antibiotic G418. Arrows indicate the position of primers 1-4 used to detect integration of the pSLI-TGD vector. Asterisks indicate a stop codon. GFP, green fluorescent protein; hDHFR; human dehydrofolate reductase for resistance to WR99210; NeoR, neomycin-resistance; 2A, Skip peptide. (B) Confirmation of vector integration for the ZNF4-KO parasites by diagnostic PCR using gDNA obtained from ZNF4-KO and WT-NF54. 5’-integration was detected using primers 1 and 4 (1164 bp) and 3’-integration using primers 3 and 2 (955 bp). Primers 3 and 4 were used to detect the presence of episome (965 bp), and primers 1 and 2 were used for WT control (1194 bp). (C) Confirmation of truncated ZNF4 tagged with GFP. Parasite lysates obtained from different stages of the ZNF4-KO parasite line were subjected to Western blotting using polyclonal mouse anti-GFP (estimated size 53 kDa). Lysate from WT mature gametocyte was used as negative control. Immunoblotting with mouse anti-Pf39 antisera (39 kDa) served as a loading control. (D) Verification of GFP expression in the ZNF4-KO parasites by live imaging. Live images of trophozoites (TZ) and gametocyte stage II (GCII) of the ZNF4-KO line detected GFP (green) in the parasite. Nuclei were counterstained with Hoechst 33342 (blue). Bar, 5 µm. (E). Asexual blood stage replication of the ZNF4-KO. Synchronized ring stage cultures of WT and ZNF4-KO with an initial parasitaemia of 0.25% were maintained in cell culture medium and the parasitaemia was followed over a time-period of 0 to 94 h via Giemsa stained smears. The data is a representation of one of two experiments performed in triplicate (mean ± SD). For the second experiment see Fig S2A. (F) Disruption of ZNF4 show no effect in gametocytaemia. Following two rounds of synchronization, a culture 5% ring stage parasites of the WT and ZNF4-KO was induced for gametocytogenesis and the next day the parasites were washed and grown with medium supplemented with 50 mM GlcNac to kill asexual blood stages for 5 day, then with normal medium till day 10 post induction. The gametocyte gametocytaemia was monitored by Giemsa stained blood smears from day 5 post induction. The result is a representative of one of two experiments (see Fig S2B for second experiment). (H). Gametocyte maturation in the ZNF4-KO. The development of gametocyte was compared between the WT and the ZNF4-KO by counting the gametocyte stages of 50 gametocytes at each time point in triplicate (See Fig 2C for second experiment). (H) Gametocyte morphology in the ZNF4-KO line. Giemsa stained pictures of gametocyte stages of the ZNF4-KO and WT.

After successful gene disruption, we first examined the asexual blood stage replication in the ZNF4-KO line. To this end, the development of highly synchronized ring stages of the ZNF4-KO and WT parasite lines was monitored over a period of 96 h by Giemsa stained blood smears, which were taken every 12 h. The results show no significant difference in intraerythrocytic replication between the ZNF4-KO and the WT (Fig 2E). Then rings stage parasites after the second replication cycle were induced for gametocytogenesis and asexual blood stages were eliminated. Gametocyte development and gametocytaemia were followed from day 5 post-activation till day 10. No significant effect in gametocytaemia and gametocyte development was observed (Fig 2F, G). In addition, the morphology of ZNF4-KO gametocytes was not affected (Fig 2H).

### Disruption of ZNF4 impacts microgamete exflagellation and reduces transmission in the mosquito

To determine if male gametocytes of the ZNF4-KO parasite line are able to produce motile microgametes, an *in vitro* exflagellation assay was performed. To this end, mature gametocytes of the ZNF4-KO and WT were activated with xanthurenic acid (XA) for 15 min at RT. After activation, the numbers of exflagellation centres were counted microscopically. The results show that the ZNF4-KO produced significantly lower numbers of exflagellation centres as compared to WT (Fig 3A) indicating an impairment in male gametocyte exflagellation. To see if this impairment can affect malaria transmission in the mosquitoes, the matured gametocytes were fed to *Anopheles stephensi* mosquito in membrane feeding assays and the number of oocysts were counted. Although the ZNF4-KO line still produced oocysts, their numbers were relatively very low as compared to the WT (Fig 3B).

**Fig 3:**
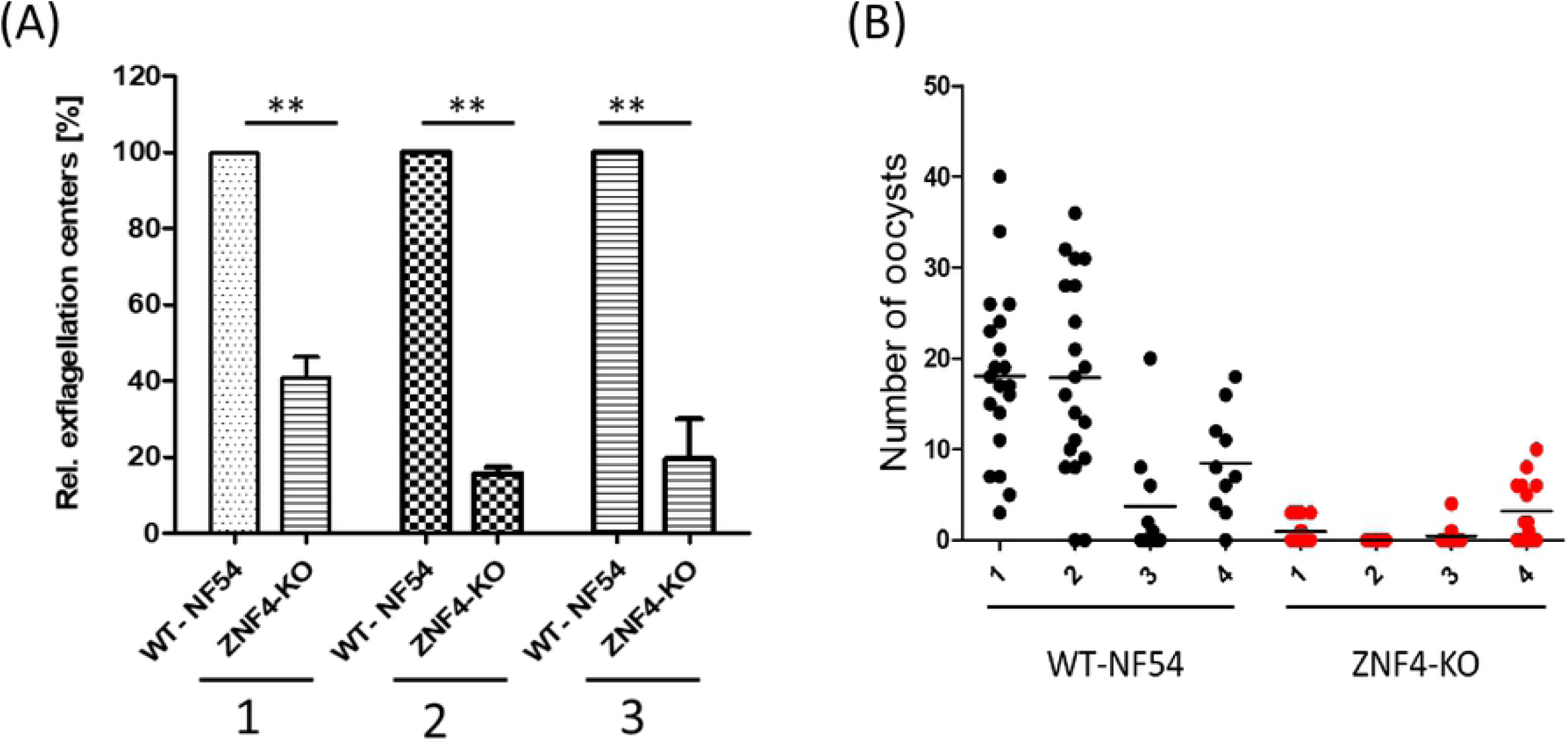
ZNF4-KO impairs male gametocyte exflagellation and parasite transmission in the mosquito. (A) Disruption of ZNF4 impairs male gametocyte exflagellation. Mature WT and ZNF4-KO gametocytes were activated *in vitro* and the number of exflagellation centres counted in 30 fields in triplicated using the light microscope. Three independent experiments were performed indicated in numbers 1 to 3. **, *P* < 0.05, Student’s *t* test. (B) Mosquito infectivity of ZNF4-KO. Enriched mature gametocytes of WT or the ZNF4-KO were fed to *An. stephensi* mosquitoes via standard membrane feeding assays. The numbers of oocysts per midgut were counted at day 10 post infection in four independent experiments each.

### ZNF4 disruption results in down-regulation of male gametocyte enriched transcripts

To compare the transcriptome of the ZNF4-KO and the WT, we carried out a comparative RNA-Seq using ring stage parasites and mature gametocytes of the ZNF4-KO and the WT. We used a cut off of greater than 2-fold change in expression. The results show that 66 genes were de-regulated in the ring stage following ZNF4 disruption (55 down-regulated and 11 up-regulated) as opposed to mature gametocytes to which a total of 473 genes (274 up-regulated and 199 down-regulated) were detected (Fig 4A, Table S1). For further analysis, we then focused on the de-regulated genes in mature gametocytes, since the knockout phenotype suggests that male gametocyte exflagellation was affected. To validate the RNA-Seq data, we carried out a quantitative real-time experiment (qRT) to compare the transcript expression of 8 genes (5 up-regulated and 3 down-regulated) using RNA from mature gametocytes of the ZNF4-KO and the WT. The results confirmed the RNA-Seq data as four of the five up-regulated genes in the RNA-Seq data were up-regulated and all down-regulated genes were down-regulated in the qRT (Fig 4B).

**Fig 4:**
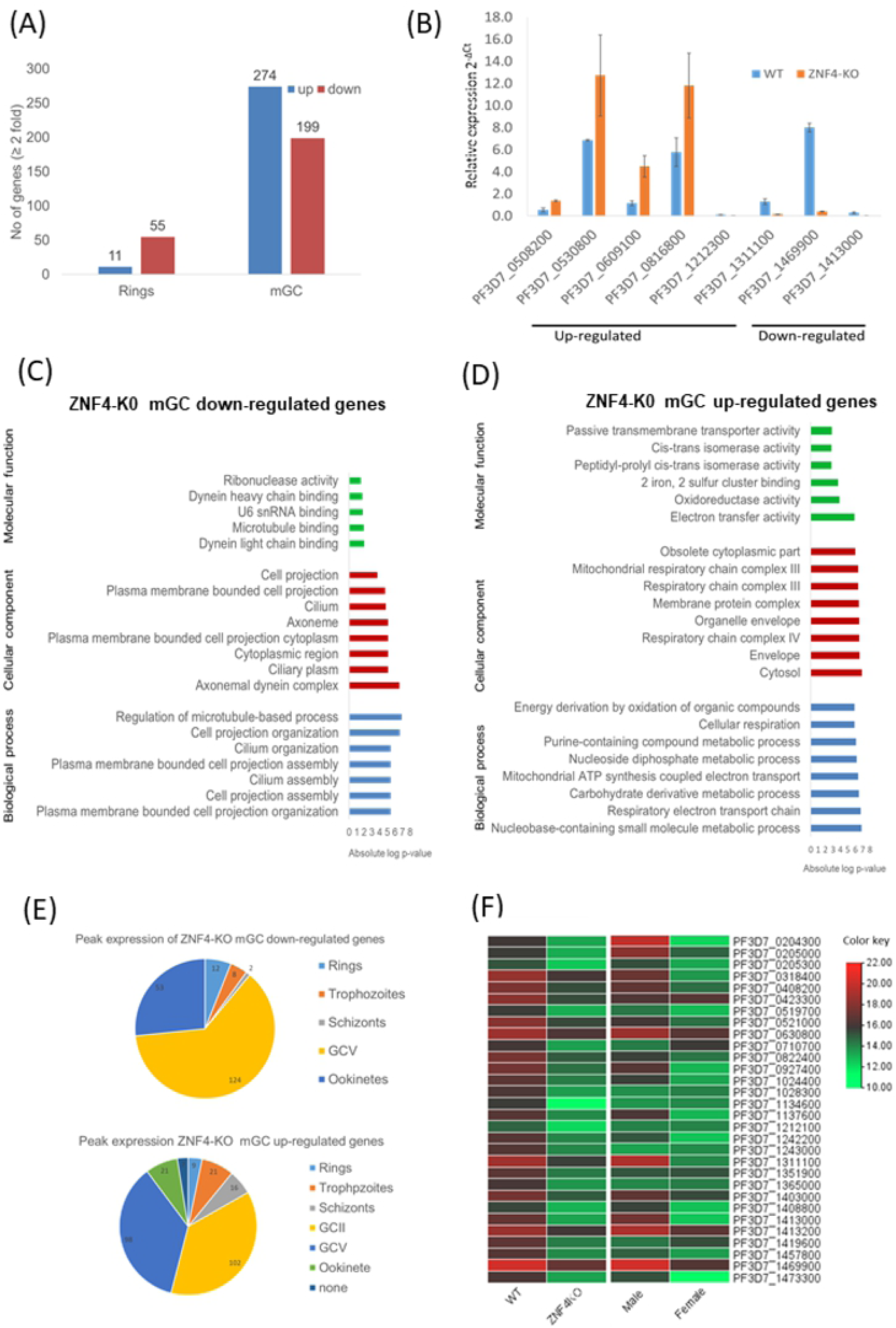
De-regulation of gene expression following disruption of ZNF4. (A) Comparative transcriptomics of de-regulated genes in the ZNF4-KO parasite line in rings and mature gametocytes. Total RNA was extracted from ring stage parasites as well as mature gametocytes of the ZNF4-KO and the WT and subjected to comparative transcriptomics by RNA sequencing. De-regulated genes with relative gene expression greater than 2-fold was considered significant. (B) Validation of RNA-Seq data by qRT. Transcript analysis for 5 up-regulated genes and 3 down-regulated in mature gametocyte as identified by RNA-Seq were validated by real-time RT-PCR. Transcript expression levels were calculated by the 2^-ΔCt^ method; the threshold cycle number (Ct) was normalized with the Ct of the gene encoding seryl tRNA-ligase (PF3D7_0717700) as reference. Genes were considered up-regulated when the changes between ZNF4-KO and WT sample were greater than 2-fold. (C, D) Summary of Gene Ontology functional analysis of mature gametocyte differentially expressed genes following ZNF4 disruption. GO enrichment analysis of de-regulated genes was determined using PlasmoDB (https://plasmodb.org/plasmo/app). The most significantly (P < 0.05) enriched GO terms in biological process, cellular component and molecular function are presented. For the complete list see Table S1. All adjusted statistically significant values of the terms were the absolute log10 values. GO, gene ontology. (E) Pie chart showing peak expression stage of de-regulated genes. The peak expression of the down-regulated as well as up-regulated genes were determined using the 7 stage RNA-Seq data [28]. (F) Heat map of top 30 down-regulated genes and their sex-specific expression. Heat map representing patterns of the top 30 down-regulated genes and their sex specific expression pattern [29]. Heat map was constructed using TB tools [30].

We next performed a gene ontology (GO) enrichment analysis (Table S1). Down-regulated genes could mainly be assigned to biological processes such as regulation of microtubule-based processes, cell projection organization and cilium organization (Fig 4C). Regarding cellular components, axonemal dynein complex assembly was highly represented and for molecular function, dynein light chain binding and microtubule binding was also highly represented (Fig 4C). Up-regulated genes were mainly assigned to nucleoside containing small molecule metabolic processes and the respiratory electron transfer chain as biological processes. In addition, the respiratory chain complex, the mitochondrial respiratory chain complex (cellular component) as well as electron transfer activity and oxidoreductase activity (molecular function) were the mostly represented (Fig 4D).

We also determined at which stage of the parasite life cycle does the de-regulated genes show peak expression according to the seven stage RNA-Seq data [28]. We observed that most of the down-regulated genes show peak expression in stage V gametocytes followed by the ookinete stage (Fig 4E). On the other hand, the up-regulated genes showed peak expression in stage II and stage V gametocytes (Fig 4E).

Since the disruption of ZNF4 mainly affected male gametocyte exflagellation, we analyzed if the effect was due to the down-regulation of transcripts enriched in male gametocytes. For this reason, we compared the sex specific transcript expression of the top 30 down-regulated genes using the sex specificity data from Lasonder and colleagues [29]. The heat map shows that indeed most of the down-regulated genes have high transcript levels in male gametocytes with genes like PF3D7_1469900 (male gametocytes enriched transcribe, MGET) and PF3D7_1311100 (meiosis-specific nuclear structural protein 1, putative) present (Fig 4F).

### ZNF4-KO up-regulated and down-regulated genes show different predicted enriched RNA binding motifs

To determine the predicted RNA-binding motif in the de-regulated mature gametocyte genes, the comprehensive motif analysis tool XSTREME (Meme-suite.org; [31]) was used to check which motifs were enriched in the up-regulated and down-regulated genes as compared to the control transcripts which were not affected. We used downloaded transcript sequences from the PlasmoDB website (plasmodb.org/plasmo/app) containing the 5’and 3’UTR and search for motif of 7 to 15 nucleotides. In the up-regulated genes motifs identified were U-rich with the top two hits being “UUUUUUUUUUUUAU” with this signature found in 199 of the 274 up-regulated transcripts (e -value: 1.8e-015; Table 1) and “AUUUUUAUUUU” with this signature found in 207 of the 274 genes (e-value: 1.5e-010; Table 1). On the other hand, motifs of the down-regulated genes were mainly A-rich with top hits being “AAAAUAUAAAAAAA” with this signature in 138 of the 199 down-regulated genes (e-value: 3.3e-012; Table 1) and “AAAAAAAAGAAAA” with signature in 165 out of the 199 down-regulated genes and (e-value: 5.8e-011; Table 1). Interestingly, these motifs were highly similar to known RBP motifs (Table 1). This indicates that ZNF4 is probably binding to different motifs in the up-regulated and down-regulated genes.

**Table 1:**
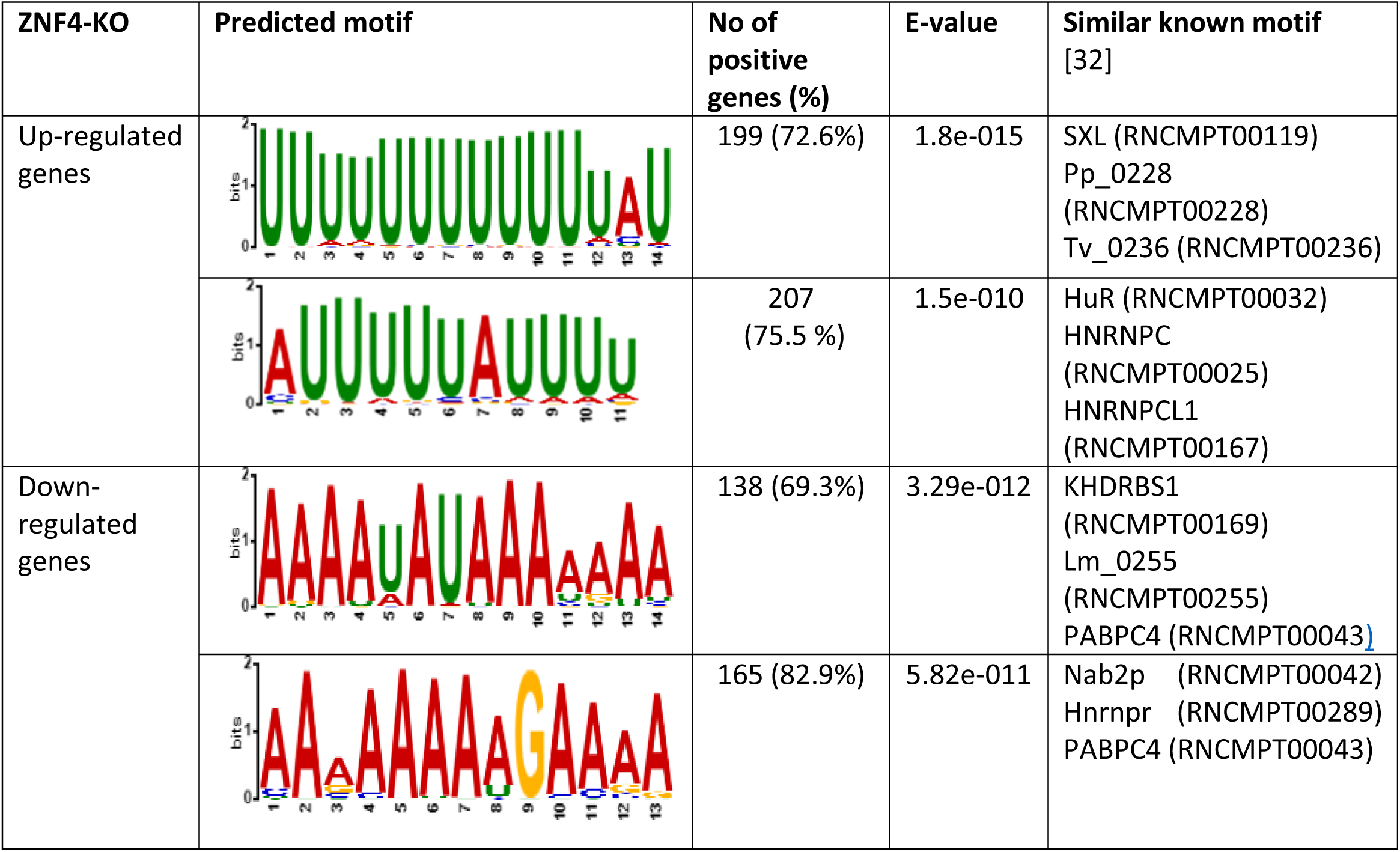
Predicted binding motif for ZNF4-KO in the up-regulated and down-regulated genes.

| ZNF4-KO | Predicted motif | No of positive genes (%) | E-value | Similar known motif [32] |
| --- | --- | --- | --- | --- |
| Up-regulated genes |  | 199 (72.6%) | 1.8e-015 | SXL (RNCMPT00119)<br>Pp_0228 (RNCMPT00228)<br>Tv_0236 (RNCMPT00236) |
|  |  | 207 (75.5 %) | 1.5e-010 | HuR (RNCMPT00032)<br>HNRNPC (RNCMPT00025)<br>HNRNPCL1 (RNCMPT00167) |
| Down-regulated genes |  | 138 (69.3%) | 3.29e-012 | KHDRBS1 (RNCMPT00169)<br>Lm_0255 (RNCMPT00255)<br>PABPC4 (RNCMPT00043) |
|  |  | 165 (82.9%) | 5.82e-011 | Nab2p (RNCMPT00042)<br>Hnrnpr (RNCMPT00289)<br>PABPC4 (RNCMPT00043) |

## Discussion

The highly complex life cycle of the human malaria parasite *P. falciparum* requires a well-coordinated gene regulation to allow for gametocyte commitment, development and human to mosquito transmission of the parasite. Translational repression in particular has been shown to be the main player in preparing female gametocytes for parasite transmission with RNA-binding translational repressors like DOZI, CITH or Puf2 playing crucial roles. While the mechanism by which female gametocytes genes are regulated has been studied in detail, the mechanism of regulation of male-specific genes remains largely unknown. In the present study, we show that the PfCCCH ZFP named ZNF4 is a potential RBP important for male gametogenesis and hence malaria transmission through the regulation of male-enriched gametocyte genes. Noteworthy, ZNF4 is expressed mainly in the cytoplasm of both male and female gametocytes in accord with previous sex-specificity data [29], indicating that ZNF4 expression is not dependent of the gametocyte sex of the parasite.

Targeted gene disruption of *ZNF4* showed normal progression through the intraerythrocytic replication cycle indicating that the gene is dispensable for asexual blood stage replication. This is not surprising as a recent genome-wide transposon mutagenesis screen in *P. falciparum* also indicates that the gene is not essential for parasite viability [33]. Also, the ZNF4-KO parasite line did not show any significant effect in gametocyte formation and development with the gametocytes showing normal morphology. However, although gametocyte development was not affected, *ZNF4* disruption greatly impaired malaria transmission in the mosquito through the inhibition of exflagellation and in consequence oocyst formation. A recent study also linked the involvement a *P. berghei* RNA binding protein UIS12 to gametocyte exflagellation and malaria transmission [34].

To determine the possible cause on exflagellation inhibition in the ZNF4-KO, we carried out comparative transcriptomic analysis in mature gametocytes as well as ring stages. Only a few genes (64) were > 2-fold de-regulated in the ring stage following *ZNF4* disruption which is in accord with the lack of any phenotype during asexual blood stage replication indicating that ZNF4 has no special role in ring stage parasites. 427 genes were de-regulated in ZNF4-KO mature gametocytes (274 genes up-regulated and 199 down-regulated), pointing to the effect in male gametocyte exflagellation.

Further analysis demonstrated that the majority of the down-regulated genes exhibit peak expression in mature gametocytes where they are implicated in essential cellular and biological processes linked to male gametogenesis such as cell projection assembly, cilium assembly and the axonemal dynein complex. The up-regulated genes mainly exhibit peak expression in stage II gametocytes with represented cellular and biological processes associated to the respiratory electron chain and mitochondrial ATP synthesis. Interestingly, a previous study which integrated transcriptomics and proteomic, associated male gametocytes to be enriched in proteins associated to formation of flagellated gametes, axoneme formation, DNA replication and chromatin organization while female gametocytes were enriched with proteins associated to protein, lipid and energy metabolism [29]. The down-regulation of genes mainly associated to cell projection assembly, cilium assembly and axonemal dynein complex formation therefore justifies the defect in male gametocyte exflagellation and studies in other organisms have associated the processes to proper flagella formation, movement and fertility [35,36]. Regarding the up-regulated genes following ZNF4-KO, it is likely that since energy metabolism is not very necessary for male gametocytes, these genes are being repressed in male gametocytes by ZNF4 and they become up-regulated following ZNF4 disruption. In accord with our findings, the top 30 down-regulated genes showed high expression in male gametocytes. Although the function of most of the male highly expressed genes are unknown, some prominent genes encode for PF3D7_1469900 (PfMGET), an abundant protein transcribed specifically in male gametocytes which has been used for male gametocyte quantification [37,38] and for PF3D7_1311100 (meiosis-specific nuclear structural protein 1, putative) with human homolog MSN1 which has been linked to male fertility [39,40].

An interesting finding in this study is the fact that up-regulated genes and down-regulated genes possess different binding motifs suggesting that they may be regulated differently. The motifs for the up-regulated genes have been reported to bind RBPs such as SXL and HuR. SXL also known as sex lethal is a master regulator of sex determinant in *Drosophilia melanogaster* by regulating the choice between male and female development pathways [41,42]. For the down-regulated gene motifs, they have been reported as targets for RBPs like Nab2p and

PABC4. Nabp2 in *Saccharomyces cerevisiae* is a nuclear protein required to protect early mRNA and has also been reported to be involved in RNA export from the nucleus to the cytoplasm [43,44]. PABC4 (cytoplasmic poly(A) binding protein C4) exhibit a critical role in erythroid differentiation through mRNA regulation [45].

The exact mechanism by which ZNF4 regulates gene expression warrants further investigation. One potential mechanism could be by translational repression as has been shown for female gametocytes where mRNA transcripts important for zygote/ookinete development such as P25 and P28 are stored in a messenger ribonucleoprotein complex composed of RNA binding protein like DOZI and CITH in *P. berghei* [14,15] or Puf2 and 7-Helix-1 in *P. falciparum* [13,46]. Noteworthy, a recent study in *P. berghei* has identified a CCCH domain containing ZFP, Pb103 to be associated to zygote/ookinete development probably by translational repression [47]. It is likely that there exists a translational repression mechanism regulating male gametocyte genes in which ZNF4 is part of the complex.

Another probable mechanism by which ZNF4 controls transcripts is by the regulation of mRNA stability as has been reported for many CCCH-ZFPs such as the tristretraprolins (TTPs) which are the most studied CCCH-ZFPs. TTPs have been shown to bind AU-rich elements in mRNAs, resulting in the removal of the poly-A tail from the mRNA, thereby marking them for decay [48]. It is possible that the deficiency of ZNF4 leads to the accumulation of transcripts due to the absence of mRNA stabilization as observed with the up-regulated genes.

In conclusion, we have identified a novel ZFP, ZNF4 and demonstrated that ZNF4 plays an essential role in the regulation of male gametocyte genes which are important of microgamete exflagellation, fertilization and parasite transmission in the mosquito. However, further studies will be required to address the mechanism by which ZNF4 regulates the gene expression and if more critical regulatory proteins are involved.

## Materials and Methods

### Antibodies

Antibodies used in this study included: rabbit anti-HA (Sigma Aldrich, Taufkirchen, Germany), rat anti-HA (Roche, Basel, Switzerland), mouse anti-GFP (Roche, Basel, Switzerland), mouse/rabbit anti-Pfs230 [49], rabbit/mouse anti-Pf39. Mouse anti-ZNF4 was generated for this study (see below). For indirect immunofluorescence assays (IFAs), the following dilutions of the antibodies were used: mouse/rabbit anti-Pfs230 (1:200), mouse anti-ZNF4 (1:20), mouse anti GFP (1:200), rabbit anti HA (1:50). For Western blot analysis the following dilutions were used: rat/rabbit anti-HA (1:500), rabbit anti-Pf39 (1:10000), anti-GFP (1:1000).

### Parasite culture

The *P. falciparum* gametocyte-producing strain NF54 was used as background strain in this study. The parasites were cultivated *in vitro* in RPMI 1640/HEPES medium (Gibco, Thermo Scientific Waltham, USA) supplemented with 10% heat-inactivated human serum and A^+^ erythrocytes at 5% hematocrit as described [50]. As supplement, 50 μg/ml hypoxanthine (Sigma Aldrich, Taufkirchen, Germany) and 10 μg/ml gentamicin (Gibco, Thermo Scientific Waltham, USA) were added to the cell culture medium and the cultures were grown in an atmosphere of of 5% O2, 5% CO2, 90% N2 at a constant temperature of 37°C. Cultures were synchronized by repeated sorbitol treatment as described [51].

Human erythrocyte concentrate and serum were purchased from the Department of Transfusion Medicine (University Hospital Aachen, Germany). The University Hospital Aachen Ethics commission approved all work with human blood, the donors remained anonymous and serum samples were pooled.

### Generation of mouse antisera

A recombinant protein, corresponding to a portion of ZNF4 (Fig 1A), was expressed as maltose-binding protein-tagged fusion protein using the pMAL(tm)c5X-vector (New England Biolabs, Ipswich, USA). The coding DNA sequence was amplified by PCR using gene-specific primers (for primer sequences, see Table S2). Recombinant protein was expressed in *E. coli* BL21 (DE3) RIL cells according to the manufacturer’s protocol (Invitrogen, Karlsruhe, Germany) and isolated and affinity-purified using amylose resin according to the manufacturer’s protocol (New England Biolabs, Ipswich, USA). Polyclonal antisera were generated by immunization of 6-weeks old female NMRI mice (Charles River Laboratories, Wilmington, USA) subcutaneously with 100 µg recombinant protein emulsified in Freund’s incomplete adjuvant (Sigma Aldrich, Taufkirchen, Germany) followed by a boost after 4 weeks. At day 10 after the boost, mice were anesthetized by intraperitoneal injection of a mixture of ketamine and xylazine according to the manufacturer’s protocol (Sigma Aldrich, Taufkirchen, Germany), and immune sera were collected via heart puncture. The immune sera of three mice immunized were pooled; sera of three non-immunized mice (NMS) were used as negative control. Experiments in mice were approved by the animal welfare committee of the District Council of Cologne, Germany (ref. no. 84-02.05.30.12.097 TVA).

### Generation of ZNF4-KO parasite line

Disruption of *ZNF4* (PF3D7_1134600) was achieved by selection-linked integration as described [27]. Briefly, the plasmid pSLI-TGD-GFP was modified to contain a 601 bp homology block from the 5’ end of the *ZNF4* coding region (for primer sequence see Table S2). Parasites were transfected as described [18] and WR99210 (Jacobus Pharmaceuticals, New Jersey, USA) was added to a final concentration of 4 nM, starting at 6 h after transfection to select integrated parasites. WR99210-resistant parasites appeared after 21 days and they were treated with medium containing 400 μg/ml G418 (Sigma Aldrich, Taufkirchen, Germany) and correct integration confirmed by diagnostic PCR (for primer sequences, see Table S2). After successful integration was obtained, the lines were maintained through selection with 4 nM WR99210.

### Generation of ZNF4-HA-glmS parasite lines

To generate a ZNF4-HA-glmS parasite line, we used selected linked integration insertion using a pSLI-HA-glmS vector (kindly provided by Dr. Ron Dzokowski, the Hebrew University of Jerusalem) in which the plasmid was modified to contain a homology block from the 3’ end of the *ZNF4* gene excluding the stop codon (for primer sequence see Table S2). Parasites were transfected and WR99210 was added to a final concentration of 4 nM, starting at 6 h after transfection to select integrated parasites. WR99210-resistant parasites appeared after 21 days and they were treated with medium containing 400 μg/ml G418 and correct integration confirmed by diagnostic PCR (for primer sequences, see Table S2).

### RNA isolation and RNA sequencing

Total RNA was isolated from ring stage and Percoll-enriched mature (stage V) gametocytes from the ZNF4-KO and NF54 WT using the Trizol reagent (Invitrogen, Karlsruhe, Germany) according to the manufacturer’s protocol. Quality of RNA samples were assessed using a ND-1000 (NanoDrop Technologies, Thermo Scientific Waltham, USA) and by agarose gel electrophoresis.

RNA sequencing was performed at the Genomic Facility of the University Clinic at the RWTH University Aachen, Germany. Briefly, the quality of the isolated total RNA samples from ring stage and mature gametocytes of the ZNF4-KO and the WT were evaluated by Tapestation 4200 (Agilent Technologies, Santa Clara, USA). The quantities were measured by Quantus Fluorometer (Promega, Manheim, Germany). Libraries were generated with TruSeq Stranded mRNA Library Preparation kit (Illumina) from high quality total RNA samples according to the manufacturer’s protocol. The generated libraries, which pass the QC check on Tapestation 4200 (Agilent Technologies, Santa Clara, USA) were sequenced on a NextSeq 500 (Illumina) with High output Kit v2.5 (150 cycles) for paired-end sequencing according to standard procedure provided by Illumina.

Data were analyzed with the NextGen pipeline, an in house-adapted pipeline embedded in the workflow management system of the QuickNGS-Environment [52]. In detail, the data were first demultiplexed according to corresponding indices. After quality assessment of the resulted fastq files with FastQC (v0.11.5), STAR v2.5.2b [53] was applied to align the reads to the *P. falciparum* 3D7 EPr1 (version 43) with default parameters. Aligned reads were quantified with Stringtie v1.3.6 as described [54]. Counts for transcripts were counted with featureCounts subread v 1.5.1 [55] and differential expression analysis were finally conducted by comparing the transcript levels in the RNA samples of the ZNF4-KO and wildtype parasites for each stage using DESeq2 -R version 3.5.1 [55]. Raw data have been submitted to the NCBI Gene Expression Omnibus (GEO; http://www.ncbi.nlm.nih.gov/geo/) under accession number GSE196298.

### Semi-quantitative RT-PCR

To determine the transcript expression of ZNF4, total RNA was isolated from rings, trophozoites, schizonts, immature, mature and 30 min post-activated gametocytes as described above. One µg of each RNA sample was used for cDNA synthesis using the SuperScript IV First-Strand Synthesis System (Invitrogen, Karlsruhe, Germany), following the manufacturer’s instructions. The synthesized cDNA was first tested by diagnostic PCR for asexual blood stage contamination using specific primers and controls without reverse transcriptase were also used to investigate potential gDNA contamination [19]. Transcript for ZNF4 (250 bp) was amplified using ZNF4 RT primers (for primer sequences, see Table S1). The following condition was used, Initial denaturation at 94°C for 2 min, followed by 25 cycles of denaturation at 94°C for 30 s, of annealing at 45°C for 30 s, and of elongation at 72°C for 30 s, and a final extension at 72°C for 2 min.

### Real-time RT-PCR

To validate the RNA -Seq data, 1 µg of total RNA from mature gametocytes from the ZNF4-KO and WT parasite line was used for cDNA synthesis using the SuperScript IV First-Strand Synthesis System following the manufacturer’s instructions (Invitrogen, Karlsruhe, Germany). The synthesized cDNA was first verified by diagnostic PCR for asexual blood stage and DNA contamination using specific primers as described [18,19]. Primers for qRT-PCR corresponding to eight up-regulated and three down-regulated genes in the ZNF4-KO were designed using the Primer 3 software (http://frodo.wi.mit.edu/primer3/) and tested in conventional PCR using DNA or cDNA to confirm primer specificity (for primer sequences, see Table S2). Real-time RT-PCR measurements were performed using the Step One Plus Real-Time Detection System (Thermo Scientific, Waltham, USA). Reactions were performed in triplicate in a total volume of 20 μl using the maxima SyBR green qPCR master mix according to manufacturer’s instructions (Thermo Scientific, Waltham, USA). Controls without template and without reverse transcriptase were included in all qRT-PCR experiments. The levels of transcript expression were calculated by the 2^-ΔCt^ method [56] using the endogenous control gene encoding the *P. falciparum* seryl tRNA-ligase (PF3D7_0717700) as reference [57,58].

### Western blotting

Asexual blood stage parasites of the WT or mutant parasite lines were obtained following tightly synchronization of cultures with 5% sorbitol [51], while gametocytes were enriched by Percoll gradient purification [59]. Parasites were released from iRBCs with 0.05% w/v saponin/PBS for 10 min at 4°C, washed with PBS and resuspended in lysis buffer (0.5% Triton X-100, 4% w/v SDS, 0.5xPBS) supplemented with protease inhibitor cocktail (Roche, Basel, Switzerland). The lysates were then resuspended in 5 x SDS-PAGE loading buffer containing 25mM DTT, heat-denatured for 10 min at 95°C, and then separated via SDS-PAGE. After the protein have been separated, they were then transferred to Hybond ECL nitrocellulose membrane (Amersham Biosciences) according to the manufacturer’s protocol. Membranes were blocked for non-specific binding by incubation in Tris-buffered saline containing 5% skimmed milk and 1% BSA, followed by incubation with the respective primary antibody at 4°C overnight. After washing, the membranes were incubated with an alkaline phosphatase-conjugated secondary antibody directed against the first antibody (Sigma Aldrich, Taufkirchen, Germany) for 1 h at RT and developed in a solution of nitroblue tetrazolium chloride (NBT) and 5-bromo-4-chloro-3-indoxyl phosphate (BCIP; Sigma Aldrich, Taufkirchen, Germany) for 5-30 min.

### Indirect immunofluorescence assay

Mixed cultures of the WT or mutant parasite lines were air-dried on glass slides and fixed for 10 min in a methanol bath at −80°C. The RBCs were membrane permeabilized to allow access to the parasites and non-specific binding sites were blocked by incubating the fixed cells in 0.01% saponin/0.5% BSA/PBS and 1% neutral serum each for 30 min at RT. Afterwards, the preparation was then incubated with the primary antibody diluted in 0.01% saponin/0.5% BSA/PBS for 2 h each at 37°C. Binding of primary antibody was visualized by incubating the preparations with Alexa Fluor 488-conjugated secondary antibody directed against the primary antibody (Thermo Fisher Scientific, Waltham, USA) diluted in 0.01% saponin/0.5% BSA/PBS for 1 h at 37°C. The different parasite stages were detected through double-labelling with stage-specific marker primary antibodies or 0.001% w/v Evans blue (Sigma Aldrich, Taufkirchen, Germany) followed by incubation with Alexa Fluor 594-conjugated secondary antibodies (Thermo Fisher Scientific (Waltham, USA) diluted in 0.01% saponin/0.5% BSA/PBS for 1 h at 37°C. Nuclei were highlighted by treatment with Hoechst nuclear stain 33342 for 10 min at RT and cells were mounted with anti-fading solution AF2 (Citifluor Ltd) and sealed with nail polish. Digital images were taken using a Leica AF 6000 microscope and processed using Adobe Photoshop CS software.

### Asexual blood stage replication assay

To compare the asexual blood stage replication between the parental WT and the ZNF4-KO line, tightly synchronized ring stage cultures were set at an initial parasitemia of 0.25% and the development of the parasite was followed by Giemsa-stained thin blood smears prepared every 12 h over a time-period of 96 h at nine different time points (0, 12, 24, 36, 48, 60, 72, 84, 96 post-seeding). The parasitaemia of each time point was determined microscopically at 1,000-fold magnification by counting the percentage of parasites in 1,000 RBCs.

### Gametocyte development assay

To determine the effect of ZNF4-KO on gametocyte development, WT and ZNF4-KO parasite lines were tightly synchronized twice in two replication cycles and set to a parasitaemia of 5% ring stage parasites. Gametocytogenesis was then induced by the addition of lysed RBCs (0.5ml of 50 % haematocrit lysed RBC in 15ml of culture medium) followed by washing of the cell the next day. The cultures were then maintained in cell culture medium supplemented with 50 mM GlcNAc (N-acetyl glucosamine) to kill the asexual blood stages for 5 days [60] and then maintained with normal cell culture medium till day 10 post induction. Samples were taken in triplicate every 24 h starting from day 5 post induction for Giemsa smear preparation. Gametocytaemia was determined per 1,000 RBCs and the gametocyte stages II-V at the different time points from 50 gametocytes were counted in triplicate. For each assay, two experiments were performed, each in triplicate.

### Exflagellation assay

To determine the effect of *ZNF4* disruption on the ability of male gametocytes to exflagellate, gametocytaemia of matured gametocytes of the WT and the ZNF4-KO parasite lines were determined. 100 µl of each gametocyte culture was activated *in vitro* with 100 µM XA for 15 min at RT. After activation, the numbers of exflagellation centres were counted at 400-fold magnification in 30 optical fields using a Leica DMLS microscope and the number of exflagellation centres were adjusted with the gametocytaemia. Exflagellation was calculated as a percentage of the number of exflagellation centres in the ZNF4-KO in relation to the number of exflagellation centres in the WT control (WT set to 100%).

### Membrane feeding assay

The effect of *ZNF4* depletion on malaria transmission was done by membrane feeding assays performed at the TropIQ Health Science, Nijmegen Netherlands through the support of the Infrastructure for the control of vector borne diseases (infravec2). Briefly, gametocyte cultures of the WT and the ZNF4-KO lines were set up and on day 16, when the gametocytes were fully matured, they were fed to female *Anopheles stephensi* mosquitoes using standard membrane feeding assays. After 7 days the midguts were dissected and the oocysts were counted following staining with mercurochrome.

### Statistical and online Analysis

Statistical analysis of significant differences in exflagellation between WT and ZNF4-KO was done using t-test with the help of the Graph Pad Prism software. ZNF4 domain structure and 3D structure was predicted using UniProt (https://www.uniprot.org/uniprot/Q8II18) and AlphaFold [25,26] respectively. Gene ontology (GO) enrichment analyses were determined using PlasmoDB (plasmodb.org/plasmo/app). For GO analysis, the default settings were used with p < 0.05. To compare transcript expression of top 30 down-regulated genes, a heat map was constructed using TB tools [30].

To determine the enriched motifs in the de-regulated genes, the comprehensive motif analysis tool XSTREME was used (meme-suite.org,[31]). Top 2 enriched motifs between 7 to 15 nucleotides in the de-regulated genes as compared to controls (48 genes with fold change 1.00) were considered. Transcripts of de-regulated genes as well as control were downloaded from the PlasmoDB (plasmodb.org/plasmo/app).

## Acknowledgements

We thank Fabian Bick for his assistance in ZNF4 protein expression and purification.

## Author Contributions

Conceptualization: CJN and GP, Funding acquisition: CJN, GP and AF, Investigation: CJN, GP, BH and AF, Methodology: CJN, BH, AF, Supervision: CJN, GP, writing -original draft: CJN, Writing, review and editing: CJN, GP and AF.

## Supporting Information captions

**Fig S1: Validation of ZNF4 protein expression using the pSLI-ZNF4-HA-glmS parasite line**. (A) Schematic depicting the single cross over homologous recombination strategy for the generation of the ZNF4-HA-glmS parasite line using the pSLI-HA-glmS vector and primer combinations for checking successful integration. (B) Diagnostic PCR to confirm vector integration using the ZNF4-HA-glmS parasite line. 5’-integration was detected using primers 5 and 8 (1543 bp) and 3’-integration using primers 7 and 6 (1253 bp). Primers 7 and 8 were used to detect the presence of episome (1348 bp), and primers 5 and 6 were used for WT control (1448 bp). (C). Confirmation of ZNF4 tagging with HA by Western blotting. Parasite lysates obtained from gametocytes of the ZNF4-HA-glmS parasite line were subjected to Western blotting using rat anti-HA. Lysates from non-infected red blood cells (niRBC) and WT mature gametocyte (WT-GC) were used as negative control. Immunoblotting with mouse anti-Pf39 antisera (39 kDa) served as a loading control. (D) Expression of ZNF4 in different blood stages of *P. falciparum* using the ZNF4-HA-glmS. Anti-rabbit HA was used to immunolabel fixed samples of trophozoites, schizonts and gametocytes (green). Asexual blood stages (trophozoites and schizonts) were visualized by labelling with mouse anti-Pf39 and gametocytes were visualized by using mouse anti-Pfs230 (red); nuclei were highlighted by Hoechst nuclear stain 33342 (blue). Bar, 5 µm.

**Fig S2: Phenotypic characterization for ZNF4-KO**. Second experiment for the Asexual development (A), Gametocytaemia (B) and gametocyte development (C) of the ZNF4-KO as compared to WT.

**Table S1: List of ZNF4-KO de-regulated genes in gametocytes and rings and complete GO-Analysis**.

**Table S2: List of primers used in the study**.

## Notes

### Competing Interest Statement

The authors have declared no competing interest.

## References

1. WHO. World malaria report 2021. [cited 13 Jan 2022]. Available: https://www.who.int/publications/i/item/9789240040496

2. Bennink S, Kiesow MJ, Pradel G. The development of malaria parasites in the mosquito midgut. Cell Microbiol. 2016;18: 905–18. doi:10.1111/cmi.12604

3. Kuehn A, Pradel G. The coming-out of malaria gametocytes. J Biomed Biotechnol. 2010;2010: 976827. doi:10.1155/2010/976827

4. Josling GA, Llinás M. Sexual development in Plasmodium parasites: knowing when it’s time to commit. Nat Rev Microbiol. 2015;13: 573–587. doi:10.1038/nrmicro3519

5. Kafsack BFC, Rovira-Graells N, Clark TG, Bancells C, Crowley VM, Campino SG, et al. A transcriptional switch underlies commitment to sexual development in malaria parasites. Nature. 2014;507: 248–52. doi:10.1038/nature12920

6. Sinha A, Hughes KR, Modrzynska KK, Otto TD, Pfander C, Dickens NJ, et al. A cascade of DNA-binding proteins for sexual commitment and development in Plasmodium. Nature. 2014;507: 253–7. doi:10.1038/nature12970

7. Shang X, Shen S, Tang J, He X, Zhao Y, Wang C, et al. A cascade of transcriptional repression determines sexual commitment and development in Plasmodium falciparum. Nucleic Acids Res. 2021;49: 9264. doi:10.1093/NAR/GKAB683

8. Yuda M, Kaneko I, Iwanaga S, Murata Y, Kato T. Female-specific gene regulation in malaria parasites by an AP2-family transcription factor. Mol Microbiol. 2020;113: 40– 51. doi:10.1111/MMI.14334

9. Li Z, Cui H, Guan J, Liu C, Yang Z, Yuan J. Plasmodium transcription repressor AP2-O3 regulates sex-specific identity of gene expression in female gametocytes. EMBO Rep. 2021;22. doi:10.15252/EMBR.202051660

10. Kaneko I, Iwanaga S, Kato T, Kobayashi I, Yuda M. Genome-Wide Identification of the Target Genes of AP2-O, a Plasmodium AP2-Family Transcription Factor. PLOS Pathog. 2015;11: e1004905. doi:10.1371/JOURNAL.PPAT.1004905

11. Modrzynska K, Pfander C, Chappell L, Yu L, Suarez C, Dundas K, et al. A Knockout Screen of ApiAP2 Genes Reveals Networks of Interacting Transcriptional Regulators Controlling the Plasmodium Life Cycle. Cell Host Microbe. 2017;21: 11–22. doi:10.1016/J.CHOM.2016.12.003

12. Bennink S, Pradel G. The molecular machinery of translational control in malaria parasites. Mol Microbiol. 2019;112: 1658–1673. doi:10.1111/mmi.14388

13. Bennink S, von Bohl A, Ngwa CJ, Henschel L, Kuehn A, Pilch N, et al. A seven-helix protein constitutes stress granules crucial for regulating translation during human-to-mosquito transmission of Plasmodium falciparum. PLoS Pathog. 2018;14. doi:10.1371/journal.ppat.1007249

14. Mair GR, Lasonder E, Garver LS, Franke-Fayard BMD, Carret CK, Wiegant JCAG, et al. Universal Features of Post-Transcriptional Gene Regulation Are Critical for Plasmodium Zygote Development. PLoS Pathog. 2010;6: e1000767. doi:10.1371/journal.ppat.1000767

15. Mair GR, Braks JAM, Garver LS, Wiegant JCAG, Hall N, Dirks RW, et al. Regulation of sexual development of Plasmodium by translational repression. Science. 2006;313: 667–669. doi:10.1126/SCIENCE.1125129

16. Miao J, Li J, Fan Q, Li X, Li X, Cui L. The Puf-family RNA-binding protein PfPuf2 regulates sexual development and sex differentiation in the malaria parasite Plasmodium falciparum. J Cell Sci. 2010;123: 1039. doi:10.1242/JCS.059824

17. Shrestha S, Li X, Ning G, Miao J, Cui L. The RNA-binding protein Puf1 functions in the maintenance of gametocytes in Plasmodium falciparum. J Cell Sci. 2016;129: 3144– 3152. doi:10.1242/JCS.186908

18. Ngwa CJ, Kiesow MJ, Papst O, Orchard LM, Filarsky M, Rosinski AN, et al. Transcriptional Profiling Defines Histone Acetylation as a Regulator of Gene Expression during Human-to-Mosquito Transmission of the Malaria Parasite Plasmodium falciparum. Front Cell Infect Microbiol. 2017;7: 320. doi:10.3389/fcimb.2017.00320

19. Ngwa CJ, Kiesow MJ, Orchard LM, Farrukh A, Llinás M, Pradel G. The g9a histone methyltransferase inhibitor BIX-01294 modulates gene expression during plasmodium falciparum gametocyte development and transmission. Int J Mol Sci. 2019;20. doi:10.3390/ijms20205087

20. Ngwa CJ, Farrukh A, Pradel G. Zinc finger proteins of Plasmodium falciparum. Cell Microbiol. 2021;23: e13387. doi:10.1111/CMI.13387

21. Hajikhezri Z, Darweesh M, Akusjärvi G, Punga T. Role of CCCH-Type Zinc Finger Proteins in Human Adenovirus Infections. Viruses. 2020;12: 1–13. doi:10.3390/v12111322

22. Taylor GA, Thompson MJ, Lai WS, Blackshear PJ. Mitogens stimulate the rapid nuclear to cytosolic translocation of tristetraprolin, a potential zinc-finger transcription factor. Mol Endocrinol. 1996;10: 140–146. doi:10.1210/MEND.10.2.8825554

23. Pomeranz MC, Hah C, Lin PC, Kang SG, Finer JJ, Blackshear PJ, et al. The Arabidopsis tandem zinc finger protein AtTZF1 traffics between the nucleus and cytoplasmic foci and binds both DNA and RNA. Plant Physiol. 2010;152: 151–165. doi:10.1104/PP.109.145656

24. Pomeranz M, Finer J, Jang JC. Putative molecular mechanisms underlying tandem CCCH zinc finger protein mediated plant growth, stress, and gene expression responses. Plant Signal Behav. 2011;6: 647–651. doi:10.4161/PSB.6.5.15105

25. Jumper J, Evans R, Pritzel A, Green T, Figurnov M, Ronneberger O, et al. Highly accurate protein structure prediction with AlphaFold. Nature. 2021;596: 583. doi:10.1038/s41586-021-03819-2

26. Varadi M, Anyango S, Deshpande M, Nair S, Natassia C, Yordanova G, et al. AlphaFold Protein Structure Database: massively expanding the structural coverage of protein-sequence space with high-accuracy models. Nucleic Acids Res. 2022;50: D439–D444. doi:10.1093/NAR/GKAB1061

27. Birnbaum J, Flemming S, Reichard N, Soares AB, Mesén-Ramírez P, Jonscher E, et al. A genetic system to study Plasmodium falciparum protein function. Nat Methods. 2017;14: 450–456. doi:10.1038/nmeth.4223

28. López-Barragán MJ, Lemieux J, Quiñones M, Williamson KC, Molina-Cruz A, Cui K, et al. Directional gene expression and antisense transcripts in sexual and asexual stages of Plasmodium falciparum. BMC Genomics. 2011;12: 587. doi:10.1186/1471-2164-12-587

29. Lasonder E, Rijpma SR, van Schaijk BCL, Hoeijmakers WAM, Kensche PR, Gresnigt MS, et al. Integrated transcriptomic and proteomic analyses of P. falciparum gametocytes: molecular insight into sex-specific processes and translational repression. Nucleic Acids Res. 2016;44: 6087–6101. doi:10.1093/nar/gkw536

30. Chen C, Chen H, Zhang Y, Thomas HR, Frank MH, He Y, et al. TBtools: An Integrative Toolkit Developed for Interactive Analyses of Big Biological Data. Mol Plant. 2020;13: 1194–1202. doi:10.1016/J.MOLP.2020.06.009

31. Grant CE, Bailey TL. XSTREME: Comprehensive motif analysis of biological sequence datasets. [cited 21 Jan 2022]. doi:10.1101/2021.09.02.458722

32. Ray D, Kazan H, Cook KB, Weirauch MT, Najafabadi HS, Li X, et al. A compendium of RNA-binding motifs for decoding gene regulation. Nature. 2013;499: 172–177. doi:10.1038/NATURE12311

33. Zhang M, Wang C, Otto TD, Oberstaller J, Liao X, Adapa SR, et al. Uncovering the essential genome of the human malaria parasitePlasmodium falciparum by saturationmutagenesis. Science. 2018;360. doi:10.1126/SCIENCE.AAP7847

34. Müller K, Silvie O, Mollenkopf HJ, Matuschewski K. Pleiotropic Roles for the Plasmodium berghei RNA Binding Protein UIS12 in Transmission and Oocyst Maturation. Front Cell Infect Microbiol. 2021;11: 1–16. doi:10.3389/fcimb.2021.624945

35. Aprea I, Raidt J, Höben IM, Loges NT, Nöthe-Menchen T, Pennekamp P, et al. Defects in the cytoplasmic assembly of axonemal dynein arms cause morphological abnormalities and dysmotility in sperm cells leading to male infertility. PLoS genetics. 2021. doi:10.1371/journal.pgen.1009306

36. Lindemann CB, Lesich KA. Flagellar and ciliary beating: The proven and the possible. J Cell Sci. 2010;123: 519–528. doi:10.1242/jcs.051326

37. Stone W, Sawa P, Lanke K, Rijpma S, Oriango R, Nyaurah M, et al. A Molecular Assay to Quantify Male and Female Plasmodium falciparum Gametocytes: Results From 2 Randomized Controlled Trials Using Primaquine for Gametocyte Clearance. 2017;216: 457–67. doi:10.1093/infdis/jix237

38. Bradley J, Stone W, Da DF, Morlais I, Dicko A, Cohuet A, et al. Predicting the likelihood and intensity of mosquito infection from sex specific plasmodium falciparum gametocyte density. Elife. 2018;7. doi:10.7554/ELIFE.34463

39. Leslie JS, Rawlins LE, Chioza BA, Olubodun OR, Salter CG, Fasham J, et al. MNS1 variant associated with situs inversus and male infertility. Eur J Hum Genet. 2020;28: 50–55. doi:10.1038/s41431-019-0489-z

40. Ta-Shma A, Hjeij R, Perles Z, Dougherty GW, Abu Zahira I, Letteboer SJF, et al. Homozygous loss-of-function mutations in MNS1 cause laterality defects and likely male infertility. PLoS Genet. 2018;14. doi:10.1371/JOURNAL.PGEN.1007602

41. Parkhurst SM, Meneely PM. Sex determination and dosage compensation: lessons from flies and worms. Science. 1994;264: 924–932. doi:10.1126/SCIENCE.8178152

42. Wang J, Bell LR. The sex-lethal amino terminus mediates cooperative interactions in RNA binding and is essential for splicing regulation. Genes Dev. 1994;8: 2072–2085. doi:10.1101/gad.8.17.2072

43. Schmid M, Olszewski P, Pelechano V, Gupta I, Steinmetz LM, Jensen TH. The Nuclear PolyA-Binding Protein Nab2p Is Essential for mRNA Production. Cell Rep. 2015;12: 128–139. doi:10.1016/J.CELREP.2015.06.008

44. Green DM, Marfatia KA, Crafton EB, Zhang X, Cheng X, Corbett AH. Nab2p is required for poly(A) RNA export in Saccharomyces cerevisiae and is regulated by arginine methylation via Hmt1p. J Biol Chem. 2002;277: 7752–7760. doi:10.1074/JBC.M110053200

45. Kini HK, Kong J, Liebhaber SA. Cytoplasmic poly(A) binding protein C4 serves a critical role in erythroid differentiation. Mol Cell Biol. 2014;34: 1300–1309. doi:10.1128/MCB.01683-13

46. Miao J, Fan Q, Parker D, Li X, Li J, Cui L. Puf mediates translation repression of transmission-blocking vaccine candidates in malaria parasites. PLoS Pathog. 2013;9: e1003268. doi:10.1371/journal.ppat.1003268

47. Hirai M, Maeta A, Mori T, Mita T. Pb103 Regulates Zygote/Ookinete Development in Plasmodium berghei via Double Zinc Finger Domains. Pathogens. 2021;10: 1536. doi:10.3390/pathogens10121536

48. Fu M, Blackshear PJ. RNA-binding proteins in immune regulation: a focus on CCCH zinc finger proteins. Nat Publ Gr. 2016 [cited 14 Jan 2022]. doi:10.1038/nri.2016.129

49. Ngwa CJ, Scheuermayer M, Mair GR, Kern S, Brügl T, Wirth CC, et al. Changes in the transcriptome of the malaria parasite Plasmodium falciparum during the initial phase of transmission from the human to the mosquito. BMC Genomics. 2013;14: 256. doi:10.1186/1471-2164-14-256

50. Ifediba T, Vanderberg JP. Complete in vitro maturation of Plasmodium falciparum gametocytes. Nature. 1981;294: 364–6. Available: http://www.ncbi.nlm.nih.gov/pubmed/7031476

51. Lambros C, Vanderberg JP. Synchronization of Plasmodium falciparum erythrocytic stages in culture. J Parasitol. 1979;65: 418–20. Available: http://www.ncbi.nlm.nih.gov/pubmed/383936

52. Wagle P, Nikolić M, Frommolt P. QuickNGS elevates Next-Generation Sequencing data analysis to a new level of automation. BMC Genomics. 2015;16. doi:10.1186/S12864-015-1695-X

53. Dobin A, Davis CA, Schlesinger F, Drenkow J, Zaleski C, Jha S, et al. STAR: ultrafast universal RNA-seq aligner. Bioinformatics. 2013;29: 15–21. doi:10.1093/BIOINFORMATICS/BTS635

54. Pertea M, Pertea GM, Antonescu CM, Chang TC, Mendell JT, Salzberg SL. StringTie enables improved reconstruction of a transcriptome from RNA-seq reads. Nat Biotechnol 2015 333. 2015;33: 290–295. doi:10.1038/nbt.3122

55. Liao Y, Smyth GK, Shi W. featureCounts: an efficient general purpose program for assigning sequence reads to genomic features. Bioinformatics. 2014;30: 923–930. doi:10.1093/BIOINFORMATICS/BTT656

56. Livak KJ, Schmittgen TD. Analysis of Relative Gene Expression Data Using Real-Time Quantitative PCR and the 2−ΔΔCT Method. Methods. 2001;25: 402–408. doi:10.1006/meth.2001.1262

57. Salanti A, Staalsoe T, Lavstsen T, Jensen ATR, Sowa MPK, Arnot DE, et al. Selective upregulation of a single distinctly structured var gene in chondroitin sulphate A-adhering Plasmodium falciparum involved in pregnancy-associated malaria. Mol Microbiol. 2003;49: 179–191. doi:10.1046/j.1365-2958.2003.03570.x

58. Wang CW, Mwakalinga SB, Sutherland CJ, Schwank S, Sharp S, Hermsen CC, et al. Identification of a major rif transcript common to gametocytes and sporozoites of Plasmodium falciparum. Malar J. 2010;9: 147. doi:10.1186/1475-2875-9-147

59. Kariuki MM, Kiaira JK, Mulaa FK, Mwangi JK, Wasunna MK, Martin SK. Plasmodium falciparum: Purification of the various gametocyte developmental stages from in vitro-cultivated parasites. Am J Trop Med Hyg. 1998;59: 505–508.

60. Fivelman QL, McRobert L, Sharp S, Taylor CJ, Saeed M, Swales CA, et al. Improved synchronous production of Plasmodium falciparum gametocytes in vitro. Mol Biochem Parasitol. 2007;154: 119–123. doi:10.1016/j.molbiopara.2007.04.008

